# Optimizing sparse and skew hashing: faster *k*-mer dictionaries

**DOI:** 10.64898/2026.01.21.700884

**Authors:** Giulio Ermanno Pibiri, Rob Patro

**Affiliations:** DAIS, Ca’ Foscari University of Venice, Italy; Dept. of Computer Science, University of Maryland, College Park, MD 20440, USA

**Keywords:** *k*-mers, minimizers, hashing, associative data structure

## Abstract

**Motivation:** Representing a set of *k*-mers — strings of length *k* — in small space under fast lookup queries is a fundamental requirement for several applications in Bioinformatics. A data structure based on *sparse and skew hashing* (SSHash) was recently proposed for this purpose [Pibiri, 2022]: it combines good space effectiveness with fast lookup and streaming queries. It is also *order-preserving*, i.e., consecutive *k*-mers (sharing a prefix-suffix overlap of length *k*–1) are assigned consecutive hash codes which helps compressing satellite data typically associated with *k*-mers, like abundances and color sets in colored De Bruijn graphs.

**Results:** We study the problem of accelerating queries under the sparse and skew hashing indexing paradigm, without compromising its space effectiveness. We propose a refined data structure with less complex lookups and fewer cache misses. We give a simpler and faster algorithm for streaming lookup queries. Compared to indexes with similar capabilities and based on the Burrows-Wheeler transform, like SBWT and FMSI, SSHash is significantly faster to build and query. SSHash is competitive in space with the fast (and default) modality of SBWT when both *k*-mer strands are indexed. While larger than FMSI, it is also more than one order of magnitude faster to query.

**Availability and Implementation:** The SSHash software is available at https://github.com/jermp/sshash, and also distributed via Bioconda. A benchmark of data structures for *k*-mer sets is available at https://github.com/jermp/kmer_sets_benchmark. The datasets used in this article are described and available at https://zenodo.org/records/17582116.

## 1. Introduction

Efficient representation and indexing of large sets of *k*-mers (substrings of fixed length *k* appearing in longer strings) is a central primitive in modern Bioinformatics. To tackle this problem, Pibiri [2022] recently introduced a data structure based on *sparse and skew hashing* (`SSHash`, hereafter). Apart from being compact and fast to query, its distinctive feature is that it is *order-preserving*: *k*-mers that overlap by *k* — 1 characters are likely assigned *consecutive* hash codes. This property makes it particularly well suited for compressing satellite data associated with *k*-mers, such as abundances and color sets in colored De Bruijn graphs, because values referring to consecutive *k*-mers tend to be very similar (if not identical), and storing them consecutively significantly improves compression, as well as, locality of reference. Indeed, state-of-the-art indexes for colored [Fan et al., 2024; Campanelli et al., 2024] and positional [Fan et al., 2023; Patro et al., 2025] De Bruijn graphs rely on SSHash for this reason. Improving it thus directly impacts on the performance of such indexes.

In this work, we revisit the sparse and skew hashing paradigm with the goal of improving query performance without compromising space effectiveness. We introduce a range of refinements, including: 1. a less complex query logic, 2. a new layout that reduces the number of cache misses per query and leads to consistently better runtime in practice, 3. a simpler and faster algorithm for *streaming* lookup queries. This is a query modality where lookup queries are issued for consecutive *k*-mers coming from reads or longer sequences. This modality is the one typically used in practice, e.g., to perform pseudo alignment on colored De Bruijn graphs [Bingmann et al., 2019; Fan et al., 2024; Alanko et al., 2023b]. (The Supplementary Material contains a description of the construction algorithm that uses multi-threading and external-memory scaling.)

We experimentally evaluate the new `SSHash` design against the latest state-of-the-art *k*-mer dictionaries that offer similar functionalities, such as `SBWT` [Alanko et al., 2023a] and `FMSI` [Sladký et al., 2025], both based on the Burrows and Wheeler [1994] transform (BWT). Our results indicate that `SSHash` achieves significantly better query performance and faster construction times. In terms of space usage, `SSHash` is competitive with `SBWT` in the bidirectional model (i.e., when both strands of each *k*-mer are indexed). This is the *de-facto* model used by applications built atop these indexes. The space usage of `SSHash` is actually better for larger *k*. It remains, on the other hand, more space-consuming than `FMSI`, but more than an order-of-magnitude faster to query.

### Paper organization

The rest of the paper is organized as follows. Section 2 fixes notation and gives the necessary background. Section 3 describes a general indexing framework based on few, well-defined, modular components: `SSHash`, described in Section 4, is a specific instantiation of this framework as well as other indexes based on hashing. This lays the foundation for the new development in subsequent sections. Section 5 and Section 6 describe refinements to `SSHash` for random lookup queries, while Section 7 for streaming lookup queries. We experimentally compare `SSHash` against state-of-the-art dictionaries in Section 8 and conclude in Section 9.

## 2. Preliminaries

In this section we fix notation and illustrate the basic tools we use throughout the paper to solve the *k*-mer dictionary problem.

### Problem 1

(The *k*-mer dictionary problem) Given a collection 𝒳 of strings and an integer *k >* 0, build a data structure that represents all *n* distinct *k*-mers of 𝒳 so that the following queries are efficient:

- For any *k*-mer *x*, Lookup(*x*) returns a handle *h* ∈ [1..*U*] (where *U* ≥ *n*) if *x* appears in 𝒳, or ⊥ otherwise. For any *x* ≠ *y*, Lookup(*x*) ≠ Lookup(*y*) must hold.
- For any *h* ∈ [1..*U*], Access(*h*) retrieves the *k*-mer *x* if Lookup(*x*) = *h*, or returns the empty string *ε* otherwise.
- For any string *s* of length at least *k*, Lookup(*s*) answers Lookup(*x*) for all *k*-mers *x* of *s*.

### Basic notation

Given a string *s*, we indicate with |*s*| its length and with *s*[*i*..*j*) its substring of length *j* — *i* beginning with the symbol in position *i* and terminating with that in position *j* — 1, for any 1 ≤ *i < j* ≤ |*s*| + 1. (We use 1-based indexing throughout the paper.) Notation “*x* ∈ *s*” means that *x* appears as a substring of *s* and we let pos(*x, s*) be the start position of *x* in *s*.

In this paper we consider strings over the DNA alphabet {`A`, `C`, `G`, `T` }. Each symbol in this alphabet has a complement: the complement of `A` is `T` (and vice versa), and the complement of `C` is `G` (and vice versa). The *reverse complement* of the string *s*, indicated with 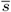, is the string where all the characters of *s* appear in reverse order and complemented. Since 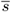 can be obtained from *s* (and vice versa), we consider them identical.

The *k*-mer *spectrum* of the string *s*, spect_*k*_ (*s*), is the set of *k*-mers of *s*. Similarly, we define spect_*k*_ (𝒳) = U_*s*∈𝒳_ spect_*k*_ (*s*). A *spectrum-preserving string set* (or SPSS) for 𝒳 is another set of strings 𝒮 = {*s*_*i*_} such that |*s*_*i*_| ≥ *k* for each *s*_*i*_ and spect_*k*_ (𝒮) = spect_*k*_ (𝒳). In this work we only consider SPSSs that do *not* repeat *k*-mers, that is spect_*k*_ (*s*_*i*_) ∩ spect_*k*_ (*s*_*j*_) = *ø* for any *s*_*i*_, *s*_*j*_ ∈ 𝒮, *i* ≠ *j*. For the rest of the paper, let 𝒮 = {*s*_*i*_} be an SPSS for 𝒳 with *N* = ∑ |*s*_*i*_|.^1^ Given a *k*-mer *x* of 𝒳, it follows that ∃! *s*_*i*_ ∈ 𝒮 such that either *x* ∈ *s*_*i*_ or 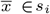. Intuitively, 𝒮 provides a natural basis for a data structure solving Problem 1 because it contains *n* distinct *k*-mers from the input 𝒳, without repetitions.

We call *S*[1..*N*] the string obtained by concatenating all the strings *s*_*i*_ in some fixed order, and indicate via *p*_*i*_, the position at which *s*_*i*_ begins in *S*. Thus, *s*_*i*_ = *S*[*p*_*i*_..*p*_*i*+1_) for *i* = 1..|𝒮| and *p*_|𝒮|+1_ = *N* + 1. Let *P* = {*p*_*i*_} in the following.

### Minimizers and super-*k*-mers

A minimizer sampling scheme is defined by a triple (*m, k*, 𝒪), where *m, k* ∈ ℕ, *m < k*, and 𝒪 is an order over all *m*-long strings. Given a *k*-mer *x*, the minimizer *µ* of *x* is the leftmost *m*-mer of *x* such that 𝒪(*µ*) ≤ 𝒪(*y*) for any other *m*-mer *y* of *x*. To simplify notation, we indicate the minimizer of the *k*-mer *x* with Mini(*x*) in the following, without specifying the parameters *m* and 𝒪. In practice, the order 𝒪 is usually random (e.g., implemented as a pseudo-random hash function).

A string is said to be *sampled* at the positions of the minimizers of its *k*-mers. For a string composed of *N* i.i.d. random characters, and when *m* is sufficiently long, the expected number of *distinct* sampled positions is approximately 2*/*(*k* — *m* + 2) *·* (*N* — *m* + 1) (see Theorem 3 by Zheng et al. [2020] for details). In other words, random minimizers are *sparse* along the string; in expectation, the start positions of two consecutive minimizers are (*k* — *m* + 2)*/*2 characters apart.

We give two more definitions. For any *k*-mer *x*, we define its *canonical minimizer* as 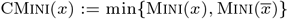

A *super-k-mer* of a string *s* is a maximal substring of *s* where a given property *R*(*x*) does not change for all *k*-mers *x* ∈ *s*. We say that *s* is *parsed* into super-*k*-mers by property *R*. Figure 1 (at page 4) illustrates an example where super-*k*-mers are parsed using (a) *R* = Mini and (b) *R* = CMini.

**Fig. 1.**
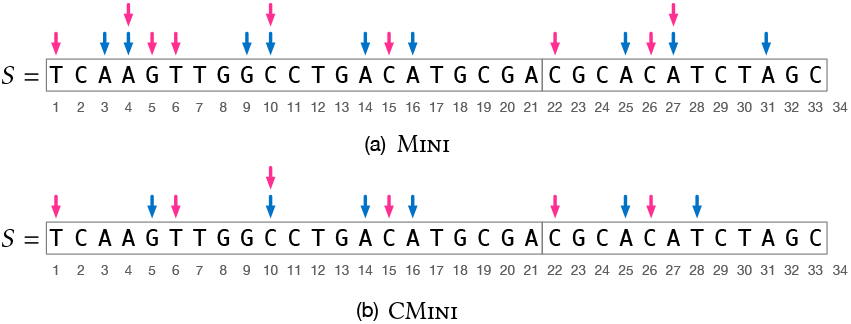
An example string S of length N = 33 resulting from the concatenation of two strings whose start positions in S are p_1_ = 1 and p_2_ = 22. There are 21 k-mers for k = 7 in the string. The blue arrows indicate the start positions of random minimizers of length m = 3, while the red arrows indicate the start positions of the corresponding super-k-mers. In (a), S is parsed into super-k-mers by Mini; in (b), by CMini. The lexicographic order of m-mers is used to compute minimizers.

### Model of computation: cache misses

To analyze the time complexity of an algorithm, we count the number of cache misses that it spends, as practical performance critically relates to them. We use the following simple model: ⌈ (*Q*) :=)*Q/B* ⌉ is the number of cache misses that occur when the algorithm reads *Q consecutive* bits, using a cache with page (or *line*) size equal to *B* bits [Vitter, 2001]. For most architectures the value *B* is 512 (64 bytes). That is, *B* spans several memory words. We assume that ⌈log_2_(*N*) ⌉ ≤ *B* for the rest of the paper^2^.

### Minimal perfect hash functions

A function *f* : *X* → {1,…, |*X*|} is said to be a minimal perfect hash function for the set *X* if *f*(*x*) ≠ *f*(*y*) for all *x, y* ∈ *X, x* ≠ *y*. In other words, a MPHF for *X* maps its keys into the first |*X*| natural numbers without collisions. Note that *f* is *not* defined for a key not in *X*, thus permitting to implement *f* with just a constant amount of space per key. Indeed the theoretical space lower bound for representing a MPHF is (less than) log_2_(*e*) ≈ 1.443 bits per key [Mehlhorn, 1982], assuming the keys of *X* are drawn from a large universe and |*X*| is large as well.

Many practical constructions for MPHFs are known that scale well to large datasets, take little space on top of the lower bound, and retain very fast evaluation time. See the recent survey by Lehmann et al. [2025] for an overview of various techniques. In this paper, we use techniques based on *bucket placement* that specialize in very fast evaluation time, requiring (under proper tuning) 1 cache miss per evaluation [Pibiri and Trani, 2021; Hermann et al., 2024; Groot Koerkamp, 2025].

### Elias-Fano sequences

Consider a sorted integer array *A*[1..*n*] such that *A*[*n*] ≤ *U* and *n* ≥ 2. The query Successor(*x*) returns the leftmost integer *y* ∈ *A* that is *y* ≥ *x* (assuming *x* ≤ *A*[*n*] w.l.o.g., so that the result is well-defined). We use the following result, based on Elias-Fano codes [Elias, 1974; Fano, 1971] (and point the interested reader to the Supplementary Material for details).

#### Theorem 1

(Elias-Fano) There exists a representation of *A* that takes at most EF(*n, U*) = *n*⌊ log_2_(*U/n*) ⌋ + 3*n* bits and:

1. For *o*(*n*) extra bits, access to the *i*-th element, can be supported with, at most, 1 + *C*_sel_ cache misses.
2. For *o*(*n*) extra bits, Successor can be supported with, at most, *C*_sel_ + *C*_scan_ cache misses.
3. For *O*(*n* log *n*) extra bits, Successor can be supported with, at most, 1 + *C*_scan_ cache misses.

The quantities *C*_sel_ and *C*_scan_ are:

- *C*_sel_ = 2 + *C*(*O*(log^4^ *n*)),
- *C*_scan_ = *C*(*U/n*) + *C*(*ℓ ·* (*U/n* + 1)) + *C*(Δ_*A*_),

Where 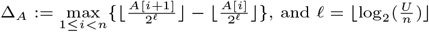.

#### Algorithm 1

*The* Lookup *algorithm for a k-mer x, using the framework illustrated in Section 3*.

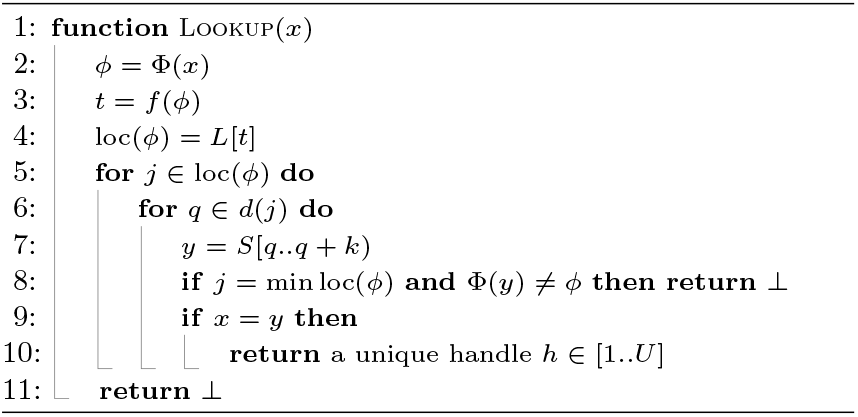

## 3. A general indexing framework

With the background and notation fixed in Section 2, we now illustrate a general indexing framework for 𝒮 that comprises four modular components. Different time/space trade-offs can be obtained depending on the choice of each component.

1. A *k*-mer transformation function Φ. This function takes a *k*-mer *x* as input and computes a value *ϕ*= Φ(*x*).
2. A locate table *L*[1..*M*], for some *M* ≤ *n*.
3. A function *f* mapping to an integer *t* ∈ [1..*M*].
4. The string *S*[1..*N*] paired with the start positions *P* = {*p*_*i*_}.

The interaction between these components is described by

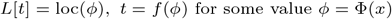

where loc(*ϕ*) is the *locate set* of *ϕ*, a sorted set of integers for which the following property holds.

### Property 1

For each *j* ∈ loc(*ϕ*), there exists a string *s*_*i*_ = *S*[*p*_*i*_..*p*_*i*+1_) and a *k*-mer *x* ∈ *s*_*i*_ such that Φ(*x*) = *ϕ* and pos(*x, S*) ∈ *d*(*j*) ⊆ [*p*_*i*_..*p*_*i*+1_ — *k*].

The function *d*(*j*) is a *displacement* function that maps the integer *j* to a set of *k*-mer positions (we will add more parameters to the function whenever necessary).

This framework is general enough to accommodate different definitions of Φ, *f*, and loc(*ϕ*), as discussed in the following with some examples.

### Queries

Algorithm 1 shows how Lookup(*x*) is implemented in the framework: the idea is to use the transformed value *ϕ* = Φ(*x*) to retrieve the set loc(*ϕ*) and use it to locate the *k*-mer *x* in *S*. Note that all *k*-mers *y* read at Line 7 must be such that Φ(*y*) = *ϕ* to be present in the input because positions in loc(*ϕ*) must satisfy Property 1. If, instead, Φ(*y*) ≠ *ϕ*, it means that the value *ϕ* was not observed for any *k*-mer in the input, hence the search can be terminated directly.

The algorithm for Access(*h*), instead, depends on the choice of *h* at Line 10 of Algorithm 1. For example, if the choice *h* = *q* is made, then *U* = *N* —*k* ≥ *n* (i.e., Lookup does not map *k*-mers into the minimal range [1..*n*], unless |𝒮| = 1) and Access(*h*) is trivial as the *k*-mer *S*[*h*..*h* + *k*) is returned directly.

Instead, if we opt for *h* = *q* — (*i* — 1) *·* (*k* — 1) where *i* is such that *p*_*i*_ ≤ *q* ≤ *p*_*i*+1_—*k*, then we make the mapping minimal, i.e., *h* ∈ [1..*n*]. In this case, the value *i* can be computed using the query Successor(*q*) over *P* (for example, representing *P* with the data structure from Theorem 1). Upon Access, however, one has to compute *i* from *h* and the sequence *P* to derive *q*, and lastly retrieve *S*[*q*..*q* + *k*).

As the number of cache misses of Algorithm 1 depends on specific choices of data structures for the framework, we defer this analysis to subsequent sections.

### Examples

Let us discuss some examples to illustrate the generality of the framework. A naïve solution would be to let Φ be a (pseudo) random hash function. Then loc(*ϕ*) would contain the positions *j* in *S* of all the *k*-mers *x* such that *ϕ* = Φ(*x*) (the displacement function would just be the identity, i.e., *d*(*j*) = {*j*}.) The key issue of this solution is, obviously, its space usage. First, *L* stores a position for each *k*-mer in the input, thus spending log_2_(*N*) bits per *k*-mer; second, the number of entries of *L* (i.e. *M*) should be chosen large enough, say *M* = Θ(*n*), to achieve fast lookup times.

This issue can be mitigated by letting *f* be a MPHF for the set of hash codes {Φ(*x*)} (assuming these are all distinct). That is, {Φ(*x*)} is mapped bijectively onto the minimal range [1..*n*] for *n* input *k*-mers, so that *M* = *n* and loc(*ϕ*) contains the unique position in *S* of the *k*-mer *x* such that *ϕ* = Φ(*x*). To save even more space, loc(*ϕ*) can store the largest multiple of a chosen quantum *v >* 1 that is smaller than (or equal to) the position of *x*, effectively spending log_2_(*N/v*) bits per *k*-mer instead of log_2_(*N*) bits. The displacement function *d* would then indicate the suitable range (of length *v*, at most). This solution is adopted by the Pufferfish index [Almodaresi et al., 2018].

Minimizers can be used to improve space usage by letting Φ = Mini and the strings in 𝒮 be parsed into super-*k*-mers by Φ. The super-*k*-mers can then be grouped by minimizer, i.e., all super-*k*-mers having the same minimizer are concatenated in the string *S*. Grouping by minimizer naturally leads to a collection of MPHFs, one function per group, to implement the mapping function *f*. Each MPHF now maps a *k*-mer appearing in the super-*k*-mers of the group to its *relative* position in the group. As positions are relative to a group, the space usage improves compared to that of Pufferfish. This is the solution implemented in the Blight index [Marchet et al., 2021].

## 4. Sparse and skew hashing

We review and analyze the sparse and skew hashing scheme (`SSHash`) [Pibiri, 2022] in this section, as it lays the foundation for the new development in subsequent sections. Also, we explain how `SSHash` can be described as a specific instance of the framework from Section 3.

### Sparse hashing

The string *S* is parsed into super-*k*-mers by Φ, where Φ is either Mini or CMini, and each *j* ∈ loc(*ϕ*) is the start position of a super-*k*-mers having minimizer *ϕ* (red arrows in the example of Figure 1 at page 4). Note that this saves space compared to Blight [Marchet et al., 2021] because the trailing *k* — 1 characters of each super-*k*-mer are encoded once (i.e., the super-*k*-mers are not grouped by minimizer but left where they appear in *S*).

Let ℳ = {Φ(*x*) | *x* ∈ spect_*k*_ (*S*)}. The function *f* is a MPHF for ℳ, hence *M* = |ℳ|. This choice of *f* also produces a space saving because the MPHF is built over the minimizers rather than the *k*-mers. As minimizers are *sparse* in the string *S*, the cost of the MPHF is small (e.g., less than 0.5 bits/*k*-mer).

A super-*k*-mer comprises at most *k* — *m* + 1 *k*-mers by construction^3^. Let *j* be the starting position of a super-*k*-mer in *S*. The displacement function is therefore as follows.

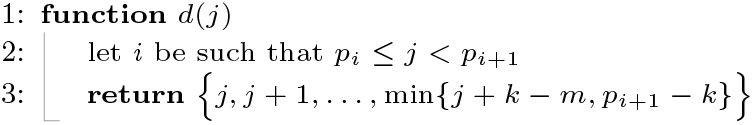

Since the index *i* is computed by *d*(*j*), the handle returned by `SSHash` at Line 10 of Algorithm 1 is *h* = *q* — (*i* — 1) *·* (*k* — 1), i.e., the *n* input *k*-mers are mapped to the minimal range [1..*n*].

### Skew hashing

Some minimizers *ϕ* may be very repetitive in *S*, hence making the set |loc(*ϕ*)| very large, and Lookup slow in the worst-case. Fortunately, using a random order and a sufficiently long minimizer length *m*, the fraction of such minimizers is very small. This is referred to as the *skew* property of minimizer occurrences. We can therefore afford to handle these few minimizers with a more space-consuming data structure that, on the other hand, guarantees a better worst-case runtime. `SSHash` adopts the following solution.

Let *l* and *r* be two integers, such that 0 ≤ *l < r*. Let *K*_*i*_ = {*x* ∈ spect_*k*_ (*S*) | 2^*i*^ *<* |loc(*ϕ*)| ≤ min{2^*i*+1^, *max*}, *ϕ*= Φ(*x*)} for *l* ≤ *i* ≤ *r*, where *max* = max_ϕ∈*M*_ |loc(*ϕ*)|. By virtue of the skew property of minimizers, we have that ∑ |*K*_*i*_| is a small fraction of the total *k*-mers. For example, we have that only ≈1.3% of the total *k*-mers belong to such sets, for the whole human genome with *k* = 31, *m* = 21, and *l* = 6. An MPHF *f*_*i*_ is built for each set *K*_*i*_. For *k*-mer *x* ∈ *K*_*i*_ with *ϕ* = Φ(*x*), we store the super-*k*-mer identifier among [1..|loc(*ϕ*)|] at position *f*_*i*_(*x*) in an array *V*_*i*_. It follows that an integer in *V*_*i*_ needs *i* +1 bits (or log_2_(*max*) bits if *i* = *r*). For the rest of the paper, we refer to a pair (*f*_*i*_, *V*_*i*_) for *l* ≤ *i* ≤ *r* as a *partition* of the skew index. We have *r* — *l* + 1 partitions (at most, when *max* ≥ 2^*r*^). At query time, if |loc(*ϕ*)| *>* 2^*l*^ then the super-*k*-mer identifier is retrieved from *f*_*i*_ and *V*_*i*_ and the query *k*-mer searched *only in one* super-*k*-mer, instead of |loc(*ϕ*)| super-*k*-mers.

### Index/query modalities

`SSHash` operates in two modalities, with different space/time trade-offs. In its *regular* modality, Φ = Mini, hence if Lookup(*x*) = ⊥ then also Lookup(*x*) is executed. In the worst-case, both loc(Mini(*x*)) and loc(Mini(*x*)) are inspected. In the *canonical* modality, Φ = CMini, so that a *k*-mer *x* and its reverse complement *x* are both mapped to the same set loc(CMini(*x*)). The Lookup algorithm is therefore executed only once (the boolean expression in the **if** at Line 9 of Algorithm 1 becomes: *x* = *y* **or**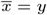) for faster query times compared to regular indexing at the price of some more space due to the higher density of canonical minimizers.

### Data structures and analysis

The string *S* is a 2-bit integer vector, as each DNA base can be coded with 2 bits. The array *P*, indicating the start position of each substring *s*_*i*_ in *S*, is coded with Elias-Fano and takes at most EF(|𝒮|,*N*) + *o*(|𝒮|) bits as per Theorem 1 (Point 1. and 2.). The locate table *L* is an array of log_2_(*N*)-bit integers, where all locate sets are stored consecutively, one after the other, in the order indicated by the MPHF *f*. Let *Z* = ∑ _*ϕ ∈ℳ*_ |loc(*ϕ*)|. The space of *L* is therefore *Z* log_2_(*N*) bits. The space for *f* is Θ(*M*) bits. With another array *sizes*[1..*M* +1] we keep track of where each loc(*ϕ*) begins and ends in *L*. That is, if *t* = *f*(*ϕ*), then loc(*ϕ*) = *L*[*sizes*[*t*]..*sizes*[*t* + 1]), with |loc(*ϕ*)| = *sizes*[*t* + 1] — *sizes*[*t*]. The sorted array *sizes* is also represented with Elias-Fano and takes at most EF(*M, Z*) + *o*(*M*) bits (Theorem 1, Point 1.). Letting *α* = ∑ |*K*_*i*_|, we upper bound the cost of the skew index by *α* log_2_(*N*)+Θ(*α*) bits, because a MPHF is built over each *K*_*i*_ and log_2_(*max*) *<* log_2_(*N*). We thus have the following result.

#### Theorem 2

(Previous `SSHash`) The space usage of `SSHash` is at most 2*N* + (*Z* + *α*) log_2_(*N*) + Θ(*M*) + Θ(*α*) + EF(|𝒮|,*N*) + EF(*M, Z*) bits. The number of cache misses per Lookup(*x*), where *z* = |loc(Mini(*x*))|, is at most

1. 1+*C*_acc_+*C*(*z* log_2_(*N*))+*z*(*C*_succ_+*C*(4*k*—2*m*)) if 1 ≤ *z* ≤ 2^*l*^,
2. 4 + *C*_acc_ + *C*_succ_ + *C*(4*k* — 2*m*) otherwise,

where *C*_acc_ = 3 + *C*(*O*(log^4^ *M*)), *C*_succ_ = 2 + *C*(*O*(log^4^ |𝒮|)) + *C*_scan_, and 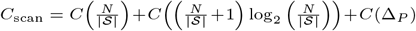.

Figure 2 in the Supplementary Material shows that the space upper bound given in Theorem 2 is quite tight. (Most of the difference between the bound and the measured space comes from overestimating the cost of the skew index.) Table 1 in the Supplementary Material, instead, compares the number of theoretical cache misses in the theorem with the measured ones in practice (and shows that they are very similar).

**Table 1.** Distinct k-mers and minimizer lengths (m) for SSHash.

| Collection | $k = 31; m$ | $k = 63; m$ |
| --- | --- | --- |
| Cod | 502,465,200; 20 | 556,585,658; 24 |
| Kestrel | 1,150,399,205; 20 | 1,155,250,667; 24 |
| Human | 2,505,678,680; 21 | 2,771,316,093; 25 |
| NCBI-v | 376,205,185; 19 | 412,515,880; 23 |
| SE | 894,310,084; 21 | 1,524,904,156; 31 |
| HPRC | 3,718,120,949; 21 | 5,926,785,469; 31 |

**Fig. 2.**
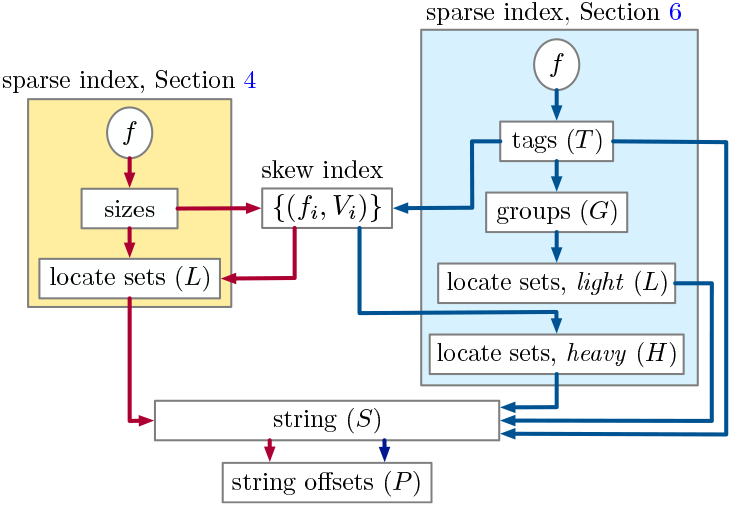
A graphical comparison between the components of the two versions of `SSHash` described in Section 4 and Section 6 (symbolic names used in the text are reported in parentheses). The skew index, S, and P, are common to both versions. In this (multi) graph, a path from f to P corresponds to the flow of execution upon Lookup (with the constraint that edges along the path do not change color). Note, in fact, that there are two possible red paths for the previous index (corresponding to the cases of Theorem 2), and three blue paths for the current one (cases of Theorem 3).

## 5. Refining displacements

The first refinement we introduce in the updated `SSHash` index regards the elements of the set loc(*ϕ*), where *ϕ* is a minimizer. As explained in Section 4, in the previous version of `SSHash`, the elements of loc(*ϕ*) are the start positions of each super-*k*-mer whose minimizer is *ϕ*. While this works seamlessly for both the regular and canonical minimizer, it also involves a linear search of a super-*k*-mer upon Lookup. To avoid the search, we now let loc(*ϕ*) := {pos(*x, S*)+pos(*ϕ, x*)— 1 | *x* ∈ spect_*k*_ (*S*) AΦ(*x*) = *ϕ*}.

### Regular displacements

At query time, given *j* ∈ loc(*ϕ*), we compute the putative start position of *x* in *S* as *q* = *j* — pos(Mini(*x*), *x*) + 1. That is, the displacement function now returns a single position, thus avoiding the need to actually search for the *k*-mer *x* in a super-*k*-mer.

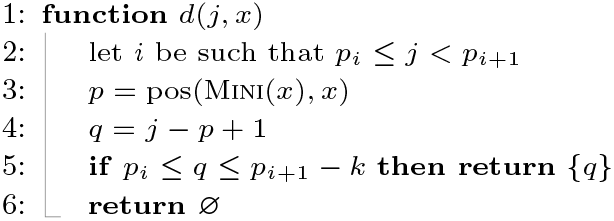

Refer to the string *S* shown in Figure 1, panel (a), and consider the query *k*-mer *x* = `AACTTGA` . As explained in Section 4, if Lookup(*x*) = ⊥ then Lookup(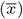) is executed. We have Mini(*x*) = `AAC` and pos(Mini(*x*), *x*) = 1. Note that no *k*-mer of *S* has minimizer `AAC`, hence *f*(`AAC`) indicates a locate set loc(*ϕ*) for another minimizer *ϕ* ≠ `AAC`, say `ATC` which begins in *S* at position *j* = 27. The position *q* = 27 — 1 + 1 = 27 is computed but *y* = *S*[27..27 + *k*) ≠ *x*. Thus, Lookup(*x*) is executed, where 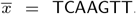. Now, 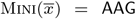 and it begins at position 3 in *x*. The candidate position *q* = 3 — 3+ 1 = 1 is computed and *x* is compared against *S*[1..1+ *k*) and produces a match.

### Canonical displacements

The canonical indexing case, where Φ = CMini, is more involved. Recall from Section 2 that we defined the canonical minimizer of a *k*-mer *x* as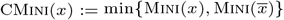. To let the new definition of loc(*ϕ*) work correctly, we also define

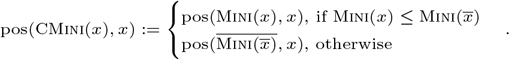

For example, consider *x* = `TCAAGTT` with 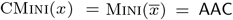. We have that pos(CMini(*x*), *x*) = 5 because 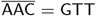 begins at position 5 in *x*.

Now, given a query *k*-mer *x*, we do not know whether it appears in *S* as *x* or as 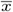 (or, if it appears at all). We therefore have to check more than one displacement per position *j* ∈ loc(CMini(*x*)) in the worst case:

1. If Mini(*x*) < 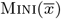): a. *j* — pos(Mini(*x*), *x*) + 1 because *x* can appear in *S*; b. 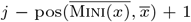 because 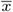 can appear in *S* and Mini(*x*) occurs in 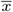 as 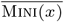.
2. If Mini(*x*) > Mini(*x*): a. 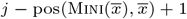 because 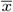 can appear in *S*; b. 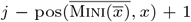 because *x* can appear in *S* and Mini(*x*) occurs in *x* as 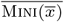.
3. 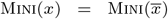 : all the four displacements are checked.

#### Observation 1

Let *ϕ* be a substring of length *m* of a *k*-mer *x*. Then 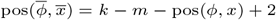.

From the previous case analysis and Observation 1, we derive the following displacement function.

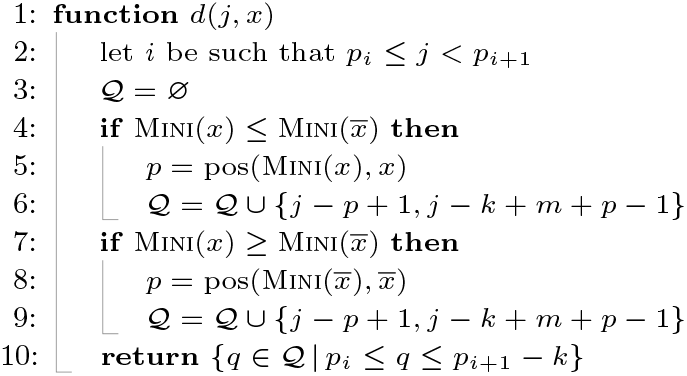

Note that for Case 1. and 2., just two displacements are computed. For Case 3., four displacements are computed. Some of them might not be valid and not returned at all (Line 10).

Consider again the string *S* in Figure 1, panel (b). Suppose we query for *x* = `TTGGCCT` . We have 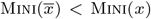 and 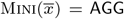, which begins at position 1 in 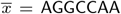. The minimizer `AGG` appears in *S* as 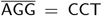 at position *j* = 10. The two positions to check, returned by *d*(*x, j*), are therefore {10, 6} and *x* is found at position 6 in *S*.

Lastly, the following observation will be useful for Theorem 3 in the analysis from Section 6.

#### Observation 2

The *k*-mers starting at positions *d*(*j, x*) occur in the same substring of *S* and this substring is at most 2*k* — *m* characters long. Hence, accessing *S* at positions in *d*(*j, x*) in ascending order costs *C*(4*k* — 2*m*) cache-misses.

## 6. Cache-efficient sparse hashing

In this section, we present a second refinement: a new data structure for sparse hashing that reduces the number of cache misses involved during Lookup.

We distinguish three *types* of minimizers based on their number of occurrences in *S*: we call a minimizer *singleton* if it appears once, *light* if it appears more than once but at most 2^*l*^ times for a small value of *l* ≥ 1, and *heavy* otherwise (it appears more than 2^*l*^ times). To reduce cache misses during Lookup queries, the idea is to reorganize the storage of the locate sets loc(*ϕ*) into three arrays according to the type of each minimizer. Let *T* [1..*M*] be an array of (log_2_(*N*) + 1)-bit integers, referred to as *tags* in the following, indexed by *f*. We use this array of tags to indicate the type of the minimizer and therefore retrieve its locate set from a dedicated array. This design allows Lookup to follow type-specific execution paths, thereby reducing cache misses. Retrieving the tag itself involves computing *f* and accessing *T* [*f*(*ϕ*)]: these two operations cost 2 cache misses.

- *Singleton minimizers*. If the minimizer appears once, the least significant bit of the tag is `0` and the remaining log_2_(*N*) bits encode the single position where the minimizer appears in *S*. At query time, whenever we read status bit `0`, we know the minimizer is singleton and the only position in the locate set is readily available in the tag without any further memory access.
- *Light minimizers*. All locate sets of size 2 ≤ *z* ≤ 2^*l*^ are stored contiguously in an array *L*, grouped by increasing size. We maintain a small auxiliary array *G*[1..2^*l*^], where *G*[*z*] stores the starting position in *L* of the group containing sets of size *z*. For a light minimizer such that *z* = |loc(*ϕ*)|, the tag consists of (from least to most significant bits): a 2-bit status field equal to `01` ; *l* bits encoding the value *z* — 2; log_2_(*N*)— *l*— 1 bits encoding the position, *i*, of loc(*ϕ*) in the group of all sets of size *z*^4^. At query time, we decode *z* and *i* from the tag and access loc(*ϕ*) = *L*[*G*[*z*]+*iz*..*G*[*z*]+(*i*+1)*z*). Retrieving *G*[*z*] costs 1 cache miss. The number of cache misses involved during a scan of loc() are *C*(|loc()| log_2_(*N*)) ≤ *C*(2^*l*^ log_2_(*N*)).
- *Heavy minimizers*. Minimizers occurring more than 2^*l*^ times are assigned status `11` and their locate sets are stored in another array *H*. These minimizers are handled with a skew index. Recall from Section 4 that a skew index comprises a collection of (at most) *r* — *l* + 1 pairs (*f*_*i*_, *V*_*i*_), where *f*_*i*_ is a MPHF and *V*_*i*_ is an vector of (*i*+1)-bit integers. Here, we choose *r* so that *r* — *l* + 1 ≤ 8 and a skew index partition identifier *i* can be coded in 3 bits. For a heavy minimizer *ϕ*, the corresponding tag stores: a 2-bit status equal to `11` ; 3 bits encoding the skew partition identifier *i*; the remaining log_2_(*N*) — 4 bits encoding the absolute offset *o* of loc(*ϕ*) in *H*. Assume *ϕ*= Φ(*x*) is a heavy minimizer. After recovering *i* and *o* from its tag, the desired position *j* ∈ loc(*ϕ*) is obtained as *j* = *H*[*o* + *V*_*i*_[*f*_*i*_(*x*)]], involving 3 cache misses.

Figure 2 illustrates how this layout differs from the previous one described in Section 4.

### Analysis

Let *β* ∈ [0, 1] be the fraction of minimizers that are not singleton (the number of singleton minimizers is (1 — *β*)*M*). The array of tags takes *M*(log_2_(*N*)+1) bits, whereas the arrays *L* and *H* take (*Z* — (1 — *β*)*M*) log_2_(*N*) bits (the array *G* is a global redundancy, which is negligible for small *l*, even if stored uncompressed as we do). The other costs are those for *S* and *f*, which are the same as those in Theorem 2, and *P* that we now represent with the data structure from Theorem 1, Point 3. Importantly, we eliminate the *sizes* array altogether. While this does not affect space too much, it is rather relevant to reduce the number of cache misses (see the next theorem). Considering the refined displacements from Section 5 and the new layout in this section, we obtain the following result.

#### Theorem 3

(Current `SSHash`) The space usage of `SSHash` is at most 2*N* + (*Z* + *α*) log_2_(*N*) + *M*(1 + *β* log_2_(*N*)) + Θ(*α*) + Θ(*M*)+*O*(|𝒮| log |𝒮|)+EF(|𝒮|,*N*)+EF(*M, Z*) bits. The number of cache misses per Lookup(*x*), where *z* = |loc(Mini(*x*))|, is at most

1. 2 + *C*_succ_ + *C*(2*k*) if *z* = 1,
2. 3 + *C*(*z* log_2_(*N*)) + *z*(*C*_succ_ + *C*(2*k*)) if 2 ≤ *z* ≤ 2^*l*^,
3. 5 + *C*_succ_ + *C*(2*k*) otherwise,

where *C*_succ_ = 1 + *C*_scan_ and *C*_scan_ is the same as that in Theorem 2. When Φ = CMini, the cost *C*(2*k*) in Case 2. becomes *C*(4*k* — 2*m*) due to Observation 2.

Assuming the same constants hidden in the Θ terms of Theorem 2 and Theorem 3, e.g., by using the same MPHF data structure, the space increase of Theorem 3 over Theorem 2 is less than *M*(*β* log_2_(*N*) — log_2_(*Z/M*) — 1) + *O*(|𝒮| log |𝒮|) bits. This space is small when *β* decreases as *m* increases. For example, *β* is on average 0.053 for *k* = 31 and *m* ≥ 19, Φ = Mini, on the datasets used in our analysis in Section 8 (see also Figure 1 and Figure 2 in the Supplementary Material.)

For the cache miss analysis in practice, refer again to Section 2 and Table 1 in the Supplementary Material.

## 7. Streaming query algorithm

Finally, we describe the simplified streaming query algorithm we adopt in the improved `SSHash` index. Let *s* be a string with |*s*| ≥ *k*. We consider the query Lookup(*s*), from Problem 1, that answers Lookup(*x*) for all *k*-mers *x* ∈ *s*.

Given that two consecutive *k*-mers of *s*, say *w* and *x*, share a (*k* — 1)-long suffix-prefix overlap, answering Lookup(*x*) *after* Lookup(*w*) should be performed faster than issuing the queries for two *k*-mers of *s* picked in any order. That is, a good implementation of Lookup(*s*) should *stream* thorough the *k*-mers of *s* to exploit the overlap information and, hence, accelerate the query.

Algorithm 2 shows the solution we use in `SSHash` . The general idea is to maintain the *state* of the last match *w* and use this information to perform faster pattern matching. For example, if the last match was found at position *q* in *S*, the next query, say for *k*-mer *x*, will compare *x* (or *x*) to *S*[*q* + 1..*q* +1 + *k*). The state of the last match *w*, is made up of several variables: its handle *h*, the identifier *i* of the comprising string, its orientation *o* ∈ {—1, +1} in *S* (under the convention that +1 indicates the forward orientation, and —1 indicates the backward orientation), whether its minimizer was *found* in *S*, its position *q* in *S*, and its minimizers Mini(*w*) and 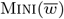. (Note that the tuple (*h, i, o, found*), computed at Line 30, can be returned by the Lookup algorithm with a straightforward modification of Algorithm 1.)

The other two variables that are part of the state are the boolean flag *start* and the integer *budget*. The *start* variable is used to let the function Minimizers (Line 14) compute correctly the minimizers of *x knowing* those of the last match *w*: that is, if *start* = **true** then *w* is not defined and the pair (Mini(*x*), 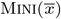) is computed from scratch; otherwise, the pair is computed in amortized *O*(1) time (using, e.g., the folklore *re-scan* method; see, the discussion in [Groot Koerkamp and Martayan, 2025; Zheng et al., 2025]). The integer *budget* defines the maximum allowable number of extensions starting from position *q* in *S*. As we extend, we decrease the budget (Line 18-21). When the budget is exhausted, we update the state of the algorithm with the function Seed.

### Algorithm 2

The Lookup(s) algorithm for a query string s. In the pseudo code, we assume an “iterator-like” interface for the string *S* such that: *S*.At(*q*) instantiates the iterator at position *q* of *S*, and *S*.Next() moves the iterator from the current position to the next and returns the *k*-mer at such position (taking into account the orientation *o* of the last match).

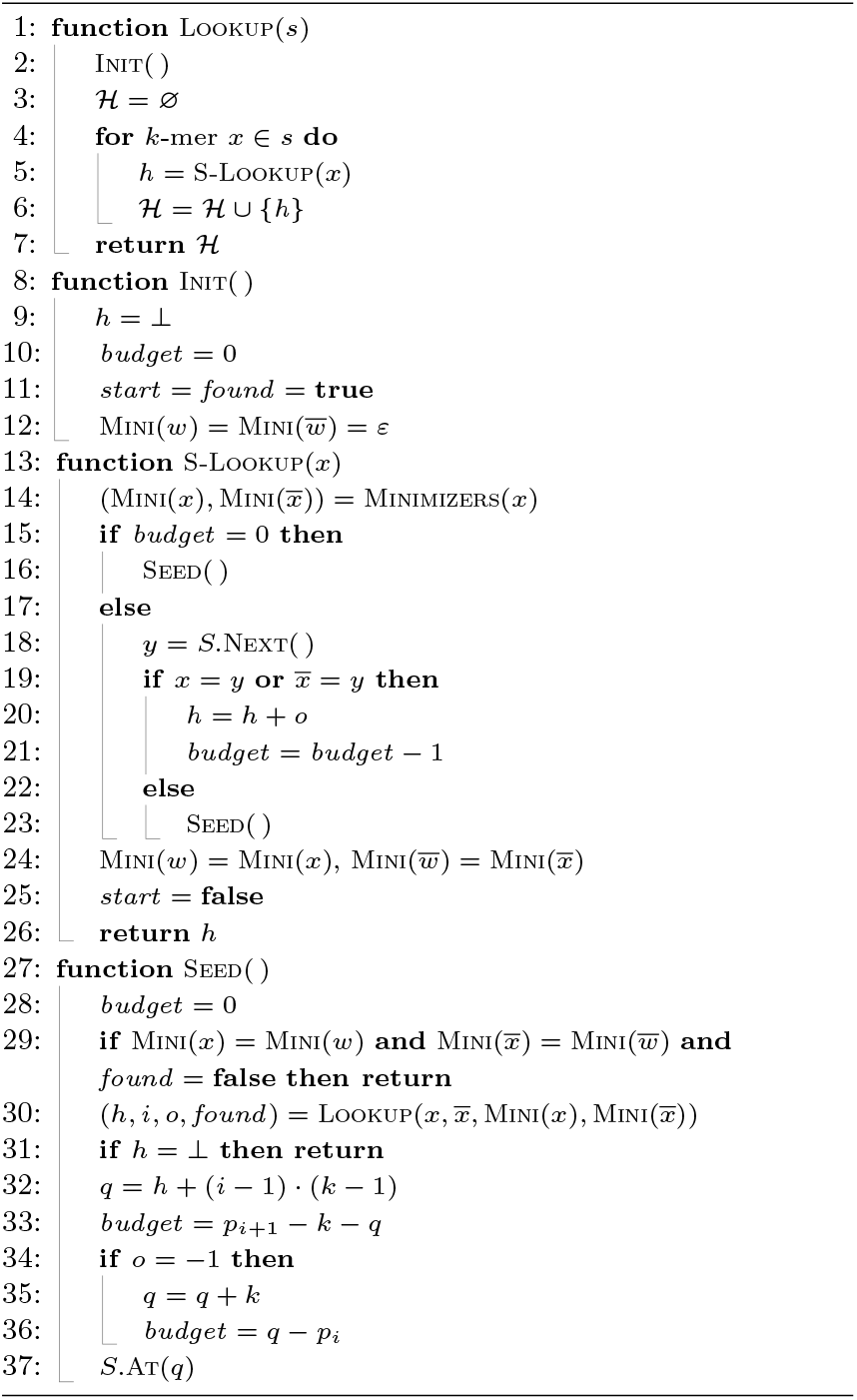

This logic is, *essentially*, the same as that described in the prior `SSHash` work (Section 4.3 of [Pibiri, 2022]) but *simpler*, thanks to the *budget* “trick”. The previous algorithm attempts to extend only if the minimizers of the last match *w* are the same as those of the current query *x*, and calls Seed whenever they change. While this already grants a fair deal of extension (because consecutive *k*-mers are likely to have the same minimizer), in the logic presented here, we attempt an extension whenever *budget* > 0 — even when minimizers change through the stream. We show in the Supplementary Material (Figure 4) that the extension rate of this refined logic is consistently higher than the previous one, granting faster query times. The two refinements presented in Section 5 and Section 6 have an important impact on the runtime of this streaming query algorithm as well.

## 8. Experimental analysis

In this section, we compare the new `SSHash` design against the two state-of-the-art *k*-mer dictionaries, `SBWT` [Alanko et al., 2023a] and `FMSI` [Sladký et al., 2025]. (The Supplementary Material also reports on the comparison against the previous published version of `SSHash` [Pibiri, 2022], using the same methodology and datasets described in this section.)

In particular, we report on the results of the experiments that were collected during November 2025 with the help of the authors of both the `SBWT` and `FMSI` . We maintain the benchmark at https://github.com/jermp/kmer_sets_benchmark, to encourage reproducibility of results. The scripts available at the repository list the precise options we used for the tools; we just report some details here.

The `SBWT` was always built by indexing both *k*-mer strands as to accelerate query processing, using the so-called “plain-matrix” variant. This is the recommended usage by the authors (and the one used in their tool Themisto [Alanko et al., 2023b] — an index for colored De Bruijn graphs based on the SBWT). Both SBWT and FMSI indexes make use of the *longest common prefix* (LCP) array to speed up queries. The space for this additional array is also included in the reported space usage.

The comparisons are shown here as bar plots for ease of visualization and space constraints, but all the data is also available in tabular format on GitHub and in the Supplementary Material.

### Hardware and compiler

All experiments were executed on a machine equipped with a AMD Ryzen Threadripper PRO 7985WX CPU, 250 GB of RAM, and a Seagate IronWolf 12 TB NAS HDD, running Ubuntu 24.04.3 LTS. All software is written in C++ and was compiled with gcc 13.3.0 using the highest optimization setting (compiler options: `-O3 -march=native` .)

### Datasets

For the experiments reported here we used two types of datasets: whole genomes and pangenomes. For the former type and for consistency with prior published work, we used the whole genomes of *Gadus morhua, Falco tinnunculus*, and *Homo sapiens* that are named Cod, Kestrel, and Human, respectively in the following. For the latter type, we used:

- NCBI-v: a collection of 18,836 virus assemblies downloaded from RefSeq in October 2025.
- SE: a pangenome containing are all the 534,751 *Salmonella enterica* genomes from Release 0.2 of the “All The Bacteria” collection [Hunt et al., 2025].
- HPRC: this is a human pangenome available (in compressed form) at https://zenodo.org/records/14854401.

From these input collections, we computed spectrum-preserving string sets in the form of eulertigs [Schmidt and Alanko, 2023] using the GGCAT algorithm [Cracco and Tomescu, 2023]. These eulertigs, along with detailed instructions about how we computed them, are available at https://zenodo.org/records/17582116, so that it is easy to reproduce our results. Table 1 reports the number of unique *k*-mers for the two values of *k* we use in this analysis, and the minimizer lengths used by `SSHash` .

### Methodology

The `SSHash` construction algorithm read the input files from the (mechanical) disk of the testing machine, and used a maximum of 16 GB of RAM and 64 threads. The same configuration was used for `SBWT`, whereas `FMSI` construction algorithm does not accept such parameters. The reported build time is the output of the Linux `time` utility.

We say Lookup(*x*) is *positive* if *k*-mer *x* is found in the dictionary (indicated with Lookup^+^ in the plots) and *negative* otherwise (indicated with Lookup^*—*^). To benchmark positive Lookup, 10^6^ *k*-mers were sampled uniformly at random from the collections and used as input. Importantly, half of them were reverse complemented to test the Lookup algorithm in the most general case. To benchmark negative Lookup instead, 10^6^ synthetic *k*-mers were created, having characters selected uniformly at random from the alphabet {A, C, G, T}. For Access, we generated 10^6^ random handles and retrieved the *k*-mers corresponding to those handles. The reported times are the averages among 5 runs of the same experiment.

To benchmark streaming Lookup queries, we take as input all the reads from experimental FASTQ files, also available at https://zenodo.org/records/17582116. These files, that contain millions of reads and are compressed with gzip, are decompressed on the fly while performing the queries. The reported time therefore also includes the incremental decompression overhead which, however, is marginal and paid by all the tested tools. The reads were chosen for each collection as to simulate a “high-hit” workload, i.e., where most *k*-mers are present in the index (say, more than 75% for *k* = 31).

All query algorithms were run using a single processing thread.

### Index space and construction efficiency

Panel (a) and (b) of Figure 3 report index space in avg. bits/*k*-mer. `FMSI` is consistently the smallest index, taking between 3.3 and 5.0 bits/*k*-mer, whereas the `SBWT` has a steady usage of 10.5 bits/*k*-mer. `SSHash` is competitive with the space usage of `SBWT`, and its space lowers substantially for larger *k* as minimizers become sparser. For this reason, in some cases like Kestrel and NCBI-v for *k* = 63, it is less than 1 bit per *k*-mer away from the effectiveness of `FMSI` .

**Fig. 3.**
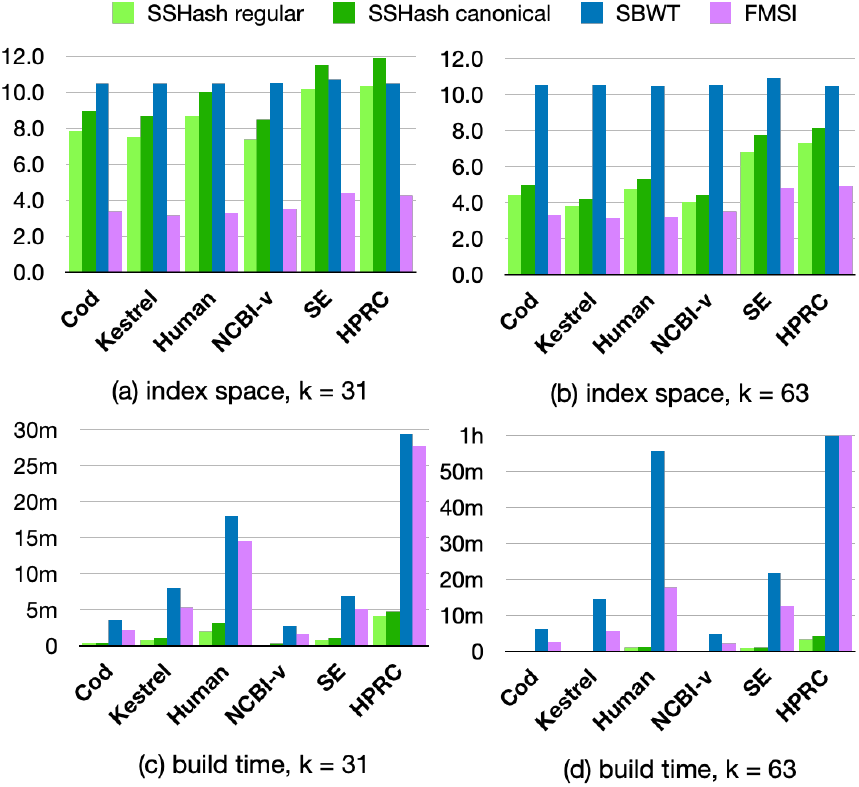
Comparison of index space in avg. bits/*k*-mer and build times. In plot (d), we cut the bars at 1h for ease of visualization: `SBWT` took 3h to build on HPRC. (`FMSI` was built within 1h.)

We consider now panel (c) and (d) of Figure 3 that report the time to build the indexes. Even on the largest collections, `SSHash` completed within 5 minutes, whereas the other tools took up to 1 hour. (For `FMSI`, we do not include the time it takes to pre-process the input to obtain the so-called *masked super-string* that it indexes.) `SBWT` is slower than `FMSI` especially for larger *k* because it enumerates and sorts *k*-mers co-lexicographically on disk (for both strands).

### Query efficiency

Figure 4 illustrates the comparison between query times. `SSHash` is generally the fastest index for all queries, by a wide margin. `FMSI` is the smallest index on disk as said before, but also the slowest to query. For example, the `SBWT` is 2× faster than `FMSI` for random Lookup queries and much faster at streaming queries. Remarkably, `SBWT` is even faster than `SSHash` on the SE dataset for streaming queries, *k* = 63. The Access query is problematic for indexes based on the BWT, as it requires tens on microseconds whereas `SSHash` takes a fraction of a microsecond.

**Fig. 4.**
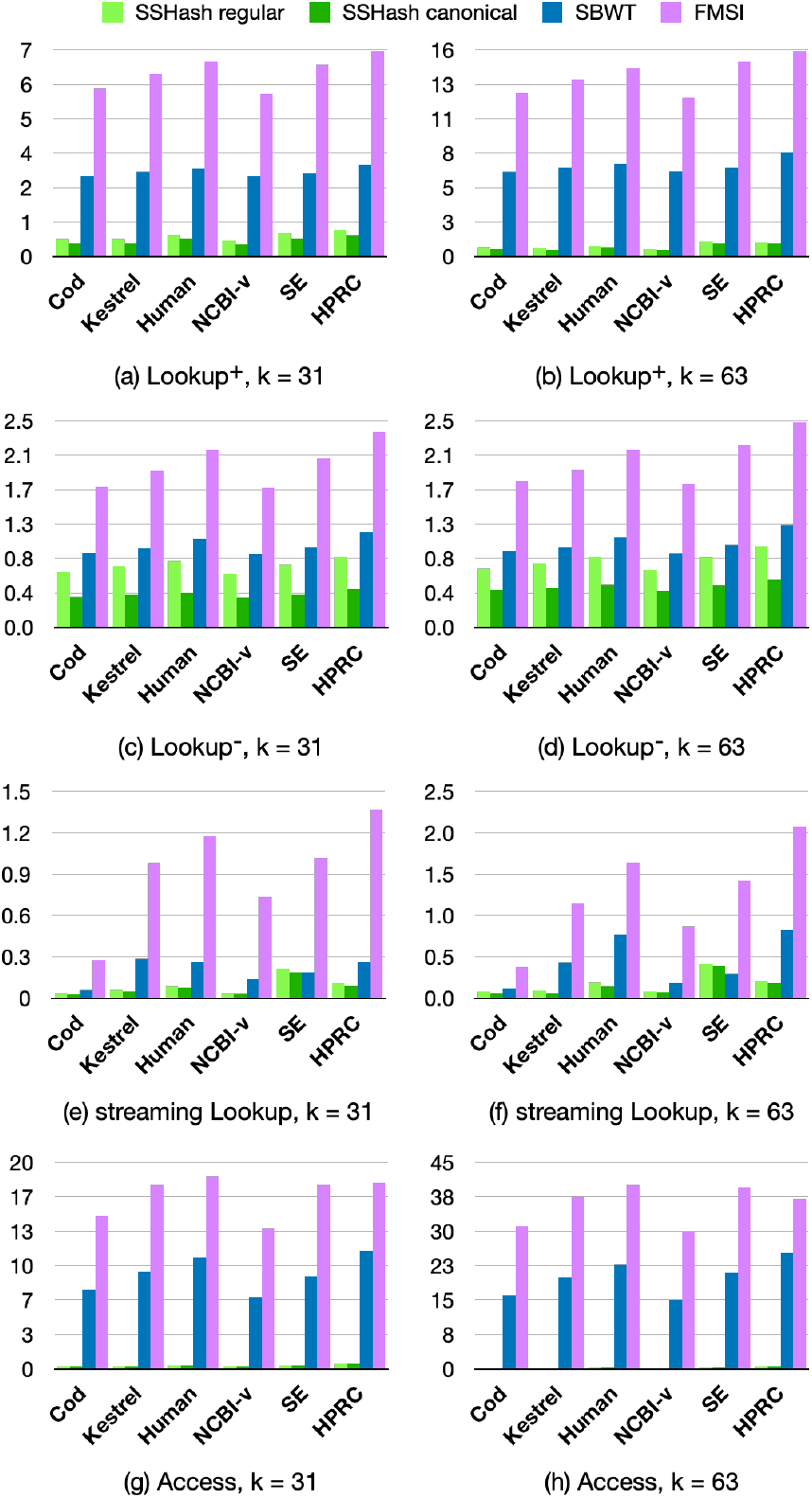
Comparison of query times in avg. *µ*s/*k*-mer.

In summary, `SSHash` regular is on average 4.2, 1.3, and 2.9 faster than the next fastest index (`SBWT`) for positive, negative, and streaming queries respectively, for *k* = 31; and 8.7, 1.2, 2.9 respectively for *k* = 63. (These factors are higher for `SSHash` canonical: 5.5, 2.5, 3.5 for *k* = 31; 10, 2, 3.6 for *k* = 63).

## 9. Conclusions and future work

In this work we improved on the sparse and skew hashing design for the *k*-mer dictionary problem. In particular, we described a simpler query procedure paired with a new layout that reduces the number of cache misses compared to the previous solution [Pibiri, 2022]. We also gave a faster streaming Lookup query algorithm. Given the fundamental nature of these queries, we expect that the speed improvements demonstrated here may lead to substantially faster downstream processing. We are genuinely surprised to see `SSHash` improve so much compared to the 2022 version and we commit ourselves to keep improving it as we critically rely on it for indexing weighted and colored De Bruijn graphs, as well as for indexing spectrum preserving tilings for reference mapping.

Compared to other indexes based on the celebrated Burrows-Wheeler transform, we found `SSHash` to be faster to query and build, but to generally consume more space. Figure 5 summarizes an example of its space/time trade-off using the whole human genome, in comparison to `SBWT` and `FMSI` .

**Fig. 5.**
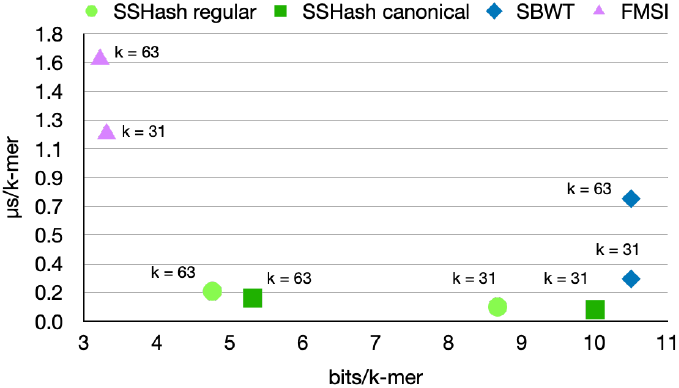
Space vs. time trade-off, for the Human dataset and streaming queries.

Future work will study the applicability of different string sampling schemes for `SSHash` . For example, preliminary experiments show that *mod-minimizers* [Groot Koerkamp and Pibiri, 2024; Groot Koerkamp et al., 2025] have the potential to reduce space consumption without hurting lookup time. Another direction could replace the locally-consistent sampling of minimizers with a *sequence-specific* one. The latter has the promise of achieving optimal density but its impact on query time, on the other hand, has yet to be analyzed.

## SUPPLEMENTARY MATERIAL

**Abstract**

Supplementary Material for the paper “Optimizing sparse and skew hashing: faster *k*-mer dictionaries”.

### 1. Elias-Fano sequences

Consider a sorted integer array *A*[1..*n*] such that *A*[*n*] ≤ *U* and *n* ≥ 2. The query Successor(*x*) returns the leftmost integer *y* ∈ *A* that is *y* ≥ *x* (assuming, w.l.o.g., *x* ≤ *A*[*n*], so that the result is well-defined). We prove the following result based on Elias-Fano codes [Elias, 1974, Fano, 1971].

#### Theorem 1

(Elias-Fano) There exists a representation of *A* that takes at most EF(*n, U*) = *n* ⌊ log_2_(*U/n*)⌋ + 3*n* bits and:

where

- *C*_sel_ = 2 + *C*(*O*(log^4^ *n*)),
- *C*_scan_ = *C*(*U/n*) + *C*(*ℓ*· (*U/n* + 1)) + *C*(Δ_*A*_),
- and 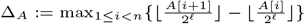,
- and *ℓ* = ⌊ log_2_(*U/n*)⌋.

#### Representation

The binary ⌈ log_2_(*U* + 1) ⌉-bit representation of each integer of *A* is split into two parts: its *ℓ* least significant bits and the remaining *h* = ⌈log_2_(*U* + 1) ⌉— *ℓ* most significant bits. We call these parts the *low* and *high* parts respectively. All the *n* low parts are written explicitly in a vector of *ℓ*-bit integers, whereas the high parts are coded using a bitvector *A*_high_ of length *n* + ⌊*U/*2^*ℓ*^⌋ + 1 bits, which is less than 3*n* bits because *n* ≤ ⌊*U/*2^*ℓ*^⌋ *<* 2*n* — 1. The main space bound, EF(*n, U*) = *n* ⌊ log_2_(*U/n*) ⌋ + 3*n*, follows.

The elements of *A* can be viewed as logically clustered into ⌊*U/*2^*ℓ*^⌋+1 clusters, 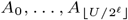, such that *A*_*j*_ contains the consecutive elements of *A* that have their high bits equal to *j*. The bitvector *A*_high_ is then

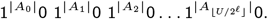

That is, it writes the cardinalities of the clusters in unary code. (Note that |*A*_*j*_ | could be 0, i.e., no element in *A* has high bits equal to *j*. In this case, the unary code is a single `0` bit. Runs of zeros might be present in *A*_high_. The length of the longest such run is Δ_*A*_.) It follows that *A*_high_ has exactly *n* bits set.

#### Random access

To decode the *i*-th value, say *A*[*i*] = *x*, from the representation, the two parts must be re-linked together. Let *x*_*ℓ*_ and *x*_*h*_ be the low and high parts of *x* respectively, so that *x* = *x*_*h*_ · 2^*ℓ*^ + *x*_*ℓ*_. The low bits *x*_*ℓ*_ are read directly from the corresponding vector, spending one cache miss. The high bits *x*_*h*_ are computed by searching *A*_high_, which can be done efficiently using Select_1_ queries. A Select_1_(*i*) query over *A*_high_ returns the position of the *i*-th one, for 1 ≤ *i* ≤ *n*. It follows that *x*_*h*_ = Select_1_(*i*) — *i*.

The extra bits used for Select_1_ as well as the number of cache misses claimed in Point 1. of Theorem 1 follow by using Theorem 2 for *A*_high_ (*u <* 3*n* and *z* = *n*). We explain Theorem 2 below.

#### Select queries

Clark [1997] shows that Select_1_ can be supported in *O*(1) time but the data structure requires a non-trivial space usage in practice and many distinct memory accesses, resulting in several cache misses.

We describe the solution by Okanohara and Sadakane [2007], the *DArray* index, which is inspired by Clark’s solution and we use in practice. (We assume Select_1_ queries throughout the presentation although one can obviously flip the ones into zeros to support Select_0_ as well.)

##### Theorem 2

(DArray) Consider a bitvector of *u* bits with *z* ones. There exists an index that takes *o*(*z*) bits and supports Select_1_ queries in at most 2 + *C*(*O*(log^4^ *u*)) cache misses.

Let *L, L*_2_, and *L*_3_ be integer quantities to be fixed later. The bitvector is split into variable-length blocks, each containing *L* ones (except for, possibly, the last block). A block is called *sparse* if its length is larger than *L*_2_, *dense* otherwise. Sparse blocks are represented verbatim, i.e., the positions of the *L* ones are coded using log_2_(*u*)-bit integers. A dense block, instead, is sparsified: we keep one `1` -bit position every *L*_3_ such positions. The positions are coded relatively to the beginning of each block, hence taking log_2_(*L*_2_) bits per position. The data structure therefore stores three arrays, *I, S, D*. The *inventory* array *I*[1..*z/L*] is such that *I*[*i*] := Select_1_(*iL*) if block *i* is dense; otherwise, *I*[*i*] = —*p* — 1 where *p* is the start position in *S* of the 1-bit positions of block *i*. The space for *I* is therefore *z/L* · log_2_(*u*) bits. The array *S* holds the positions of the *L* ones in sparse blocks. As we have at most *z/L*_2_ sparse blocks, its space is *z/L*_2_ · *L* · log_2_(*u*) bits at most. Lastly, the array *D*[1..*z/L*_3_] is such that *D*[*i*] is the position of the *iL*_3_-th one, relative to the start position of the comprising block. Its space is *z/L*_3_ · log_2_(*L*_2_) bits.

A Select_1_(*i*) query, 1 ≤ *i* ≤ *z*, first checks *p* = *I*[*i/L*]: if *p <* 0, then the block is sparse and the query is answered as *S*[−*p* − 1+ (*i* mod *L*)]; otherwise the position *D*[*i/L*_3_] is retrieved and a sequential scan of at most *L*_2_ bits of *B* is executed starting from position *p* + *D*[*i/L*_3_]. It follows that the number of cache misses per query is: 2, if *i* belongs to a sparse block; 2 + *C*(*L*_2_), if *i* belongs to a dense block.

Choosing *L* = *O*(log^2^ *u*), *L*_2_ = *O*(log^4^ *u*), and *L*_3_ = *O*(log *u*), all the three arrays *I, S*, and *D* take *o*(*z*) bits and the number of cache misses per query is at most *C*_sel_ = 2 + *C*(*O*(log^4^ *u*)). (In practice, our implementation of the Darray uses *L* = 2^10^, *L*_2_ = 2^16^, and *L*_3_ = 2^5^.)

#### Successor

Using Select_0_ queries on *A*_high_, it is also possible to support the query Successor(*x*). From *x*, we compute *x*_*h*_ = ⌊*x/*2^ℓ^ ⌋ and *i* = *p* — *x*_*h*_ with *p* = Select_0_(*x*_*h*_). For *x*_*h*_ *>* 0, this indicates that there are *i* values whose high parts are *less* than *x*_*h*_ (when *x*_*h*_ = 0, we let *i* = 0). On the other hand, *j* = Select_0_(*x*_*h*_ + 1) — *x*_*h*_ gives us the position of the first element having high bits larger than *x*_*h*_. Since a cluster contains at most 2^ℓ^ ≤ *U/n* elements, we have that *j* − *i* ≤ 2^ℓ^ ≤ *U/n* elements, and the successor could be determined by binary searching in the range *A*[*i*..*j*] for a total of *O*(log(*U/n*)(1 + *C*_sel_)) cache misses. This algorithm is not, however, cache-efficient. It is better in practice to answer the query by scanning *A* from the (*i* + 1)-th element. We follow this latter approach as it matches our implementation. (It relies on the fact that Δ_*A*_ is small for practical applications of Elias-Fano, like `SSHash`, albeit Δ = *O*(*n*) in the worst case.) When scanning from the (*i* + 1)-th element, the following cases can happen:

1. The bit in position *p* +1 of *A*_high_ is `0` : then cluster *x*_*h*_ +1 is empty and the successor of *x* is *A*[*i* + 1] (minimum element in the next non-empty cluster). The low bits of *A*[*i* + 1] are retrieved with 1 cache miss, whereas the high bits are computed by scanning *A*_high_ from position *j* until the next bit set. Since the longest run of zeros in *A*_high_ has length Δ_*A*_, *C*(Δ_*A*_) cache misses are issued during the scan.

2. The bit in position *p* + 1 of *A*_high_ is `1`, so the cluster is not empty. The elements in the cluster all have the same high bits *x*_*h*_. Now, two cases can happen:

a. The successor is not larger than the largest element in the cluster, so it belongs to the cluster. Scanning up to *U/n* elements therefore costs *C*(*U/n*)+ *C*(ℓ · *U/n*) cache misses.

b. The successor is larger than the largest element in the cluster, so it is the minimum in the next non-empty cluster. The cost is at most *C*(*U/n*)+ *C*(ℓ · (*U/n* + 1))+ *C*(Δ_*A*_).

Using again Theorem 2 on the zeros of *A*_high_ (*u <* 3*n* and *z* = 2*n*) to implement Select_0_, the extra bits and number of cache misses claimed in Point 2. of Theorem 1 follow.

Lastly, Point 3. of Theorem 1 illustrates a more space-consuming alternative that, on the other hand, supports faster Successor. The idea is to use an extra array *hints*[1.. ⌊*U/*2^*ℓ*^ ⌋ such that *hints*[*i*] = Select_0_(*i*), for *i* = 1.. ⌊*U/*2^*ℓ*^ ⌋. As *n* ≤ ⌊*U/*2^*ℓ*^ ⌋ *<* 2*n* − 1 and |*A*_high_| *<* 3*n*, the space bound follows. Instead of computing Select_0_(*x*_*h*_), this value is readily available as *hints*[*x*_*h*_] (in the general case when *x*_*h*_ *>* 0).

## 2. Cache miss analysis: theory and practice

As discussed in Section 2 of the main paper, we model the number of cache misses involved during a read of *Q* bits from main memory to the cache with *C*(*Q*) := ⌈*Q/B*⌉, where *B* is the cache line size. In this section we validate this model and show that it is accurate under proper tuning. In particular, we compare the number of theoretical cache misses of Lookup claimed in Theorem 2 and Theorem 3 of the main paper with the *actual* number of cache misses measured using the Linux <monospace>perf</monospace> tool (command: <monospace>perf stat -B -e cache-misses</monospace>).

For ease of presentation, we report again below the number of theoretical cache misses from Theorem 2 and Theorem 3 of the main paper. Both are valid for a Lookup(*x*) query, with *z* = |loc(Mini(*x*))|.

### Theorem 2

**previous SSHash**. The number of cache misses is at most

1. 1+*C*_acc_+*C*(*z* log_2_(*N*))+*z*(*C*_succ_+*C*(4*k*—2*m*)) if 1 ≤ *z* ≤ 2^*l*^;
2. 4 + *C*_acc_ + *C*_succ_ + *C*(4*k* — 2*m*) otherwise.

### Theorem 3

**current SSHash**. The number of cache misses is at most

1. 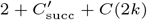
2. 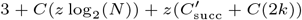
3. 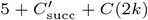

The values of *C*_acc_, *C*_succ_, 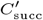, and *C*_scan_ are:

- *C*_acc_ = 3 + *C*(*O*(log^4^ *M*)),
- *C*_succ_ = 2 + *C*(*O*(log^4^ |*S*|)) + *C*_scan_,
- 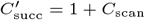
- *C*_scan_ = *C*(*N/*|𝒮|) + *C*((*N/*|𝒮| + 1) log_2_(*N/*|𝒮|)) + *C*(Δ_*P*_).

### Fixing the parameters and result

We fix *B* = 512, which is a very common cache line size and, indeed, that of our testing machine. For the choice of *k* and *m* in our experimental analysis (Section 8 of the main paper), we have *C*(2*k*) = *C*(4*k* — 2*m*) = 1 for *k* = 31 and *m < k*. Although the longest run of zeros on the high bitvector of the Elias-Fano can can be as large as *O*(*n*) in the worst case, Δ_*P*_ is actually small on tested datasets. For example, it is 140 on the whole human genome. So, we let *C*_scan_ = 3. We let *C*(*O*(log^4^ *M*)) = 6 · *C*(*O*(log^4^ |𝒮|)) for Theorem 2 because, in practice, the *sizes* array is approximately 6 times larger than the *P* array. We use *C*(*O*(log^4^ |𝒮|)) = 1 for Cod but *C*(*O*(log^4^ |𝒮|)) = 1.5 for both Human and HPRC as their respective indexes are much larger than that for Cod. For the value of *z* we use the average number of positions *j* inspected by 10^6^ random positive Lookup queries. Lastly, we have log_2_(*N*) equal to 30, 32, and 34 for Cod, Human, and HPRC respectively.

Table 1 reports the result of the comparison: for every case and dataset, the model closely matches the actual number of cache misses.

**Table 1.**
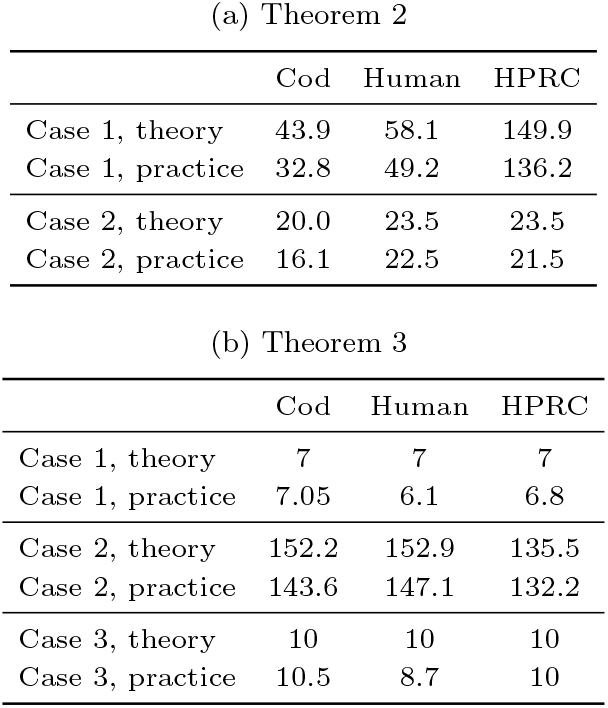
Number of cache misses: theory and practice. These results are for *k* = 31 and SSHash regular, for random positive Lookup queries.

## 3. Construction

We describe a multi-threaded construction algorithm for the new layout of `SSHash` described in Section 6 of the main paper, designed to scale to large collections using external memory and a fixed RAM budget. The construction takes as input a compressed collection of strings (in FASTA format and compressed, for example with gzip) and proceeds as a pipeline of streaming and sorting phases. In short: minimizers are first generated by streaming through the strings, in parallel; then, sorted in external memory; and finally laid out contiguously to avoid cache misses. The construction steps are as follows.

### 1. Input encoding

Each string in the input is decompressed incrementally, 2-bit encoded using SIMD instructions, and concatenated in *S*.

### 2. Parallel minimizer computation

The obtained string *S* is split into chunks, one chunk per thread. The RAM dedicated to the construction is split evenly among threads. Each thread computes the minimizers in its chunk in a *streaming* fashion.

We use the folklore *re-scan* method which performs better in practice than the monotone-queue approach we used before (see the discussion in [Zheng et al., 2025]). For each minimizer occurrence, the thread emits the tuple (ϕ, *j, p, v*) into a thread-local in-memory buffer. The tuple comprises: the minimizer itself *ϕ* (as a 2-bit encoded string), its occurrence *j* in *S*, the offset *p* indicating that the super-*k*-mer of minimizer ϕ starts in *S* at position *j* − *p* + 1, and lastly *v*, the number of *k*-mers in its super-*k*-mer. (These last two quantities, *p* and *v*, are used to build efficiently the skew index in the last step.) When the buffer reaches its dedicated capacity, the tuples in the buffer are sorted by the components (ϕ, *j*) and the buffer is flushed to disk as a sorted run.

### 3. External merge using a loser tree

The sorted runs on disk are merged into a single run using a classic multi-ary external merge algorithm. However, instead of a min-heap, a *loser tree* is used to select the minimum element at each step of the merge. A loser tree is a complete binary-tree, like a min-heap, but it performs only one comparison per tree level when updating the minimum (compared to two, as spent by a min-heap), yielding a ≈30% speedup in our experiments. Knuth [1998] (Section 5.4.1) gives a description of the loser tree.

### 4. MPHF construction

From the merged minimizer stream on disk, we build the MPHF *f* using external memory and multiple threads. Our implementation uses `PTHash` as choice of MPHF [Pibiri and Trani, 2021, 2023].

### 5. Resorting tuples in MPHF order

The minimizer tuples on disk are sorted again according to the identifier assigned to minimizers by *f* . As a result, all occurrences of the same minimizer become contiguous in a file on disk. This process is implemented, again, with a parallel external-memory merge sort.

### 6. Locate sets construction

Since minimizer tuples are now laid out consecutively on disk, the arrays *T, G, L*, and *H* are all computed by scanning the tuples sequentially.

### 7. Skew index construction

During the scan, minimizers occurring more than 2^*l*^ times are detected and the *r* − *l* + 1 partitions are built one after the other. Consider a tuple (ϕ, *j, p, v*) such that 2^*i*^ *<* |loc(ϕ)| ≤ 2^*i*+1^ for some *i* ≥ *l*. All the *k*-mers *x* ∈ *S*[*j* − *p* − 1..*j* − *p* + *v* + *k*) are added to the set *K*_*i*_ under formation. As soon as the next processed minimizer has a locate set larger than 2^*i*+1^, the MPHF *f*_*i*_ is built (in parallel) for *K*_*i*_, *V*_*i*_ laid out consequently, and the process continues with the next partition.

## 4. Experimental comparison against the previous version

We compare the version of `SSHash` from this paper (referred to as *current* in the plots) and the previous version^1^, using the same datasets, machine, and methodology described in Section 8. In general, the current version outperforms the previous one under every aspect and consistently on all tested dataset.

Figure 1 illustrates the space and build time of the two versions. The space is very similar between the two versions. Furthermore, Figure 2 shows the comparison between the actual, measured, space and the bounds from Theorem 2 and Theorem 3 in the main paper. In both cases, the bounds are quite close to the measured space and most of the difference comes from overestimating the cost of the skew index with *α*(log_2_(*N*) + Θ(1)) bits.

**Fig. 1.**
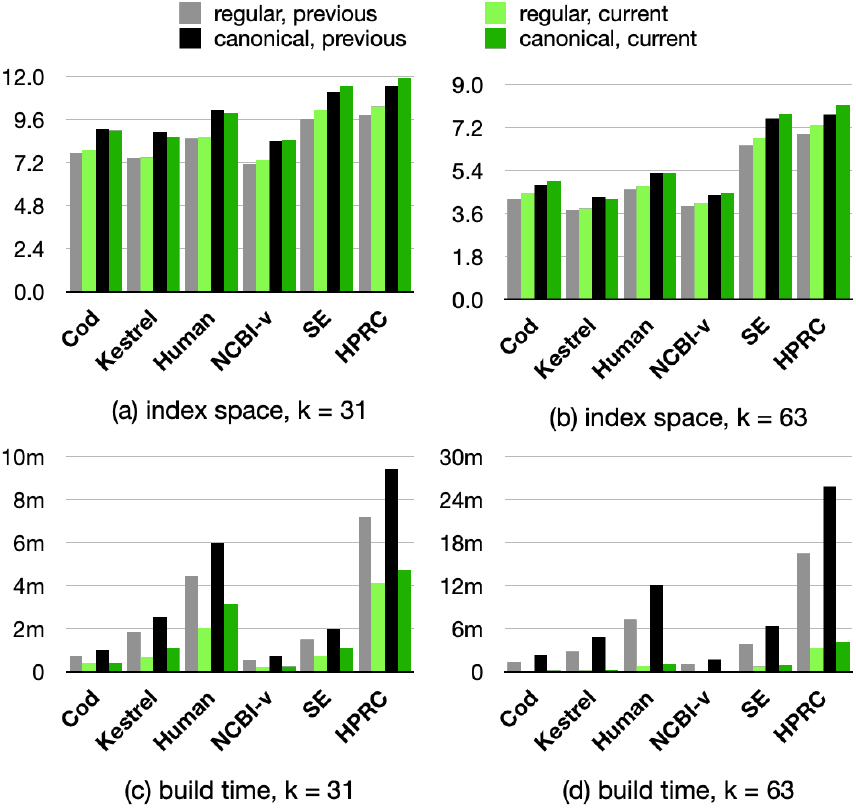
Comparison between previous and current `SSHash` : index space (in avg. bits/*k*-mer) and build time.

**Fig. 2.**
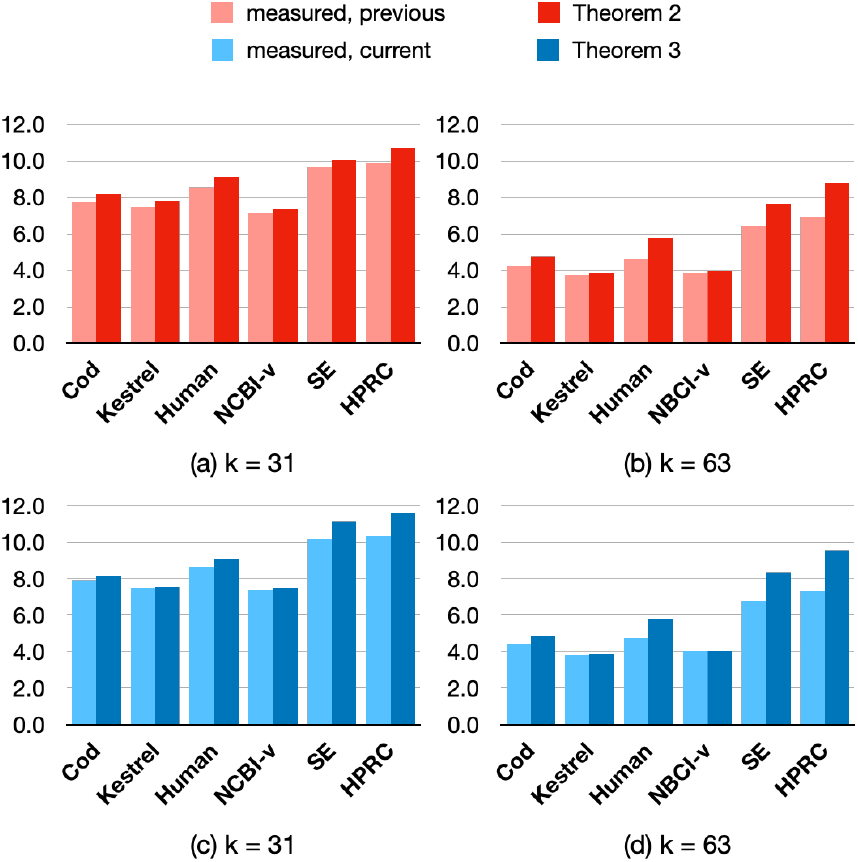
Comparison between measured index space and that from Theorem 2 and Theorem 3 of the main paper. Space is reported in avg. bits/*k*-mer. The constants in the asymptotic terms Θ(α) and Θ(*M*) are the same from both theorems and equal to 2.5 and 3.0 respectively, which are faithful to our implementation.

The current version is between 2 − 3x faster to build, on average for *k* = 31. This result improves for larger *k*; for example, it is up to 6x faster for *k* = 63. The better build time is due to better multi-threading, faster minimizer computations over streams, and faster merging in external memory which the previous version supported only partially.

Figure 3, instead, shows the query times of the two versions. (Query time for Access is almost the same between the two versions, as apparent from the tables in this document, so we do not discuss it in the following.) Avoiding the scan of super-*k*-mers and the cache-efficient layout both contribute to faster random Lookup queries. The simpler logic of the streaming Lookup algorithm paired with the more efficient random Lookup, results in 2 − 3x faster streaming queries. In particular, the refined logic consistently increases the extension rate compared to the previous version, of about 15% for *k* = 31 (Figure 4).

**Fig. 3.**
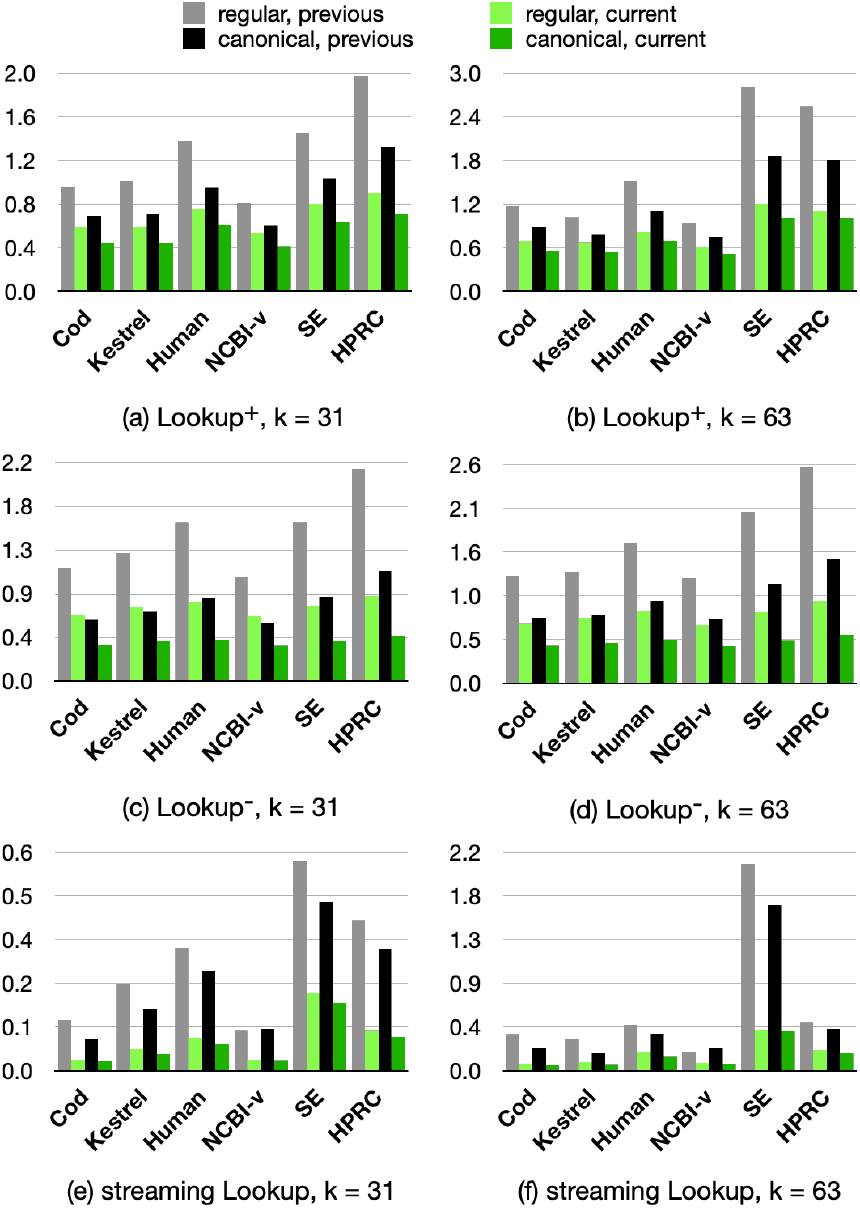
Comparison between previous and current `SSHash` : query times are reported in avg. *µ*s/*k*-mer.

**Fig. 4.**
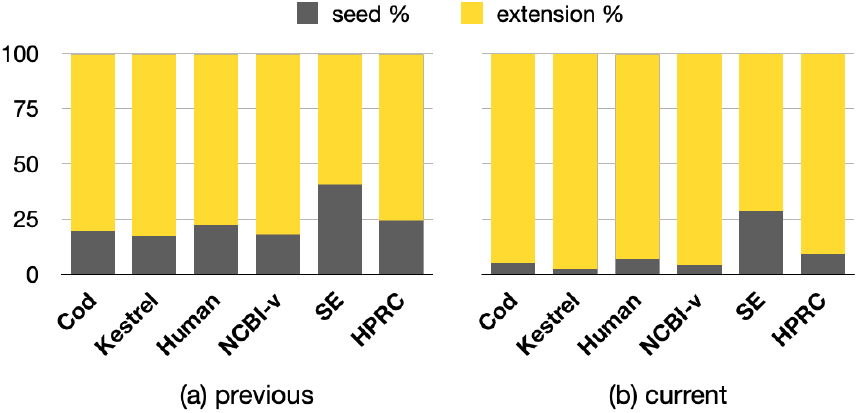
4. Seed vs. extension rate for streaming Lookup queries, for *k* = 31.

**Table 2.** Index space and construction efficiency for current SSHash.

| (a) regular |  |  |  |  |
| --- | --- | --- | --- | --- |
| $k$ | Collection | bits/$k$-mer | GB | build time |
| 31 | Cod | 7.89 | 0.50 | 26s |
|  | Kestrel | 7.50 | 1.08 | 43s |
|  | Human | 8.67 | 2.72 | 2m 1s |
|  | NCBI-v | 7.37 | 0.35 | 13s |
|  | SE | 10.17 | 1.14 | 45s |
|  | HPRC | 10.35 | 4.81 | 4m 8s |
| 63 | Cod | 4.44 | 0.31 | 12s |
|  | Kestrel | 3.82 | 0.55 | 16s |
|  | Human | 4.76 | 1.65 | 54s |
|  | NCBI-v | 4.05 | 0.21 | 5s |
|  | SE | 6.79 | 1.29 | 44s |
|  | HPRC | 7.33 | 5.43 | 3m 20s |

| (b) canonical |  |  |  |  |
| --- | --- | --- | --- | --- |
| $k$ | Collection | bits/$k$-mer | GB | build time |
| 31 | Cod | 9.01 | 0.57 | 26s |
|  | Kestrel | 8.67 | 1.25 | 1m 6s |
|  | Human | 10.01 | 3.14 | 3m 10s |
|  | NCBI-v | 8.48 | 0.40 | 16s |
|  | SE | 11.51 | 1.29 | 1m 6s |
|  | HPRC | 11.93 | 5.54 | 4m 45s |
| 63 | Cod | 4.97 | 0.35 | 15s |
|  | Kestrel | 4.22 | 0.61 | 19s |
|  | Human | 5.31 | 1.84 | 1m 9s |
|  | NCBI-v | 4.46 | 0.23 | 7s |
|  | SE | 7.77 | 1.48 | 58s |
|  | HPRC | 8.14 | 6.03 | 4m 13s |

**Table 3.** Query efficiency for current SSHash . Timings for Lookup and Access are in (average) microseconds per *k*-mer. For Streaming, we report (average) nanoseconds per *k*-mer.

| (a) regular |  |  |  |  |  |
| --- | --- | --- | --- | --- | --- |
| $k$ | Collection | Lookup<sup>+</sup> | Lookup<sup>-</sup> | Access | Streaming |
| 31 | Cod | 0.59 | 0.67 | 0.28 | 30 |
|  | Kestrel | 0.59 | 0.74 | 0.28 | 60 |
|  | Human | 0.75 | 0.80 | 0.36 | 90 |
|  | NCBI-v | 0.54 | 0.65 | 0.26 | 30 |
|  | SE | 0.80 | 0.76 | 0.36 | 213 |
|  | HPRC | 0.90 | 0.86 | 0.54 | 112 |
| 63 | Cod | 0.69 | 0.71 | 0.29 | 77 |
|  | Kestrel | 0.67 | 0.78 | 0.33 | 86 |
|  | Human | 0.82 | 0.86 | 0.36 | 188 |
|  | NCBI-v | 0.61 | 0.69 | 0.28 | 85 |
|  | SE | 1.20 | 0.85 | 0.41 | 412 |
|  | HPRC | 1.10 | 0.98 | 0.64 | 213 |

| (b) canonical |  |  |  |  |  |
| --- | --- | --- | --- | --- | --- |
| $k$ | Collection | Lookup<sup>+</sup> | Lookup<sup>-</sup> | Access | Streaming |
| 31 | Cod | 0.44 | 0.37 | 0.28 | 26 |
|  | Kestrel | 0.44 | 0.40 | 0.28 | 46 |
|  | Human | 0.61 | 0.42 | 0.35 | 74 |
|  | NCBI-v | 0.41 | 0.36 | 0.26 | 29 |
|  | SE | 0.63 | 0.40 | 0.36 | 186 |
|  | HPRC | 0.71 | 0.46 | 0.54 | 93 |
| 63 | Cod | 0.56 | 0.45 | 0.29 | 60 |
|  | Kestrel | 0.54 | 0.48 | 0.33 | 66 |
|  | Human | 0.69 | 0.52 | 0.36 | 146 |
|  | NCBI-v | 0.52 | 0.44 | 0.28 | 72 |
|  | SE | 1.00 | 0.51 | 0.41 | 400 |
|  | HPRC | 1.00 | 0.58 | 0.64 | 181 |

---

**Table 4.** Index space and construction efficiency for previous.

| (a) regular |  |  |  |  |
| --- | --- | --- | --- | --- |
| $k$ | Collection | bits/$k$-mer | GB | build time |
| 31 | Cod | 7.75 | 0.49 | 43s |
|  | Kestrel | 7.47 | 1.07 | 1m 50s |
|  | Human | 8.56 | 2.68 | 4m 27s |
|  | NCBI-v | 7.17 | 0.34 | 33s |
|  | SE | 9.65 | 1.08 | 1m 30s |
|  | HPRC | 9.88 | 4.59 | 7m 14s |
| 63 | Cod | 4.23 | 0.29 | 1m 28s |
|  | Kestrel | 3.76 | 0.54 | 2m 56s |
|  | Human | 4.63 | 1.60 | 7m 20s |
|  | NCBI-v | 3.90 | 0.20 | 1m 4s |
|  | SE | 6.46 | 1.23 | 3m 51s |
|  | HPRC | 6.94 | 5.14 | 16m 32s |
| (b) canonical |  |  |  |  |
| $k$ | Collection | bits/$k$-mer | GB | build time |
| 31 | Cod | 9.11 | 0.57 | 1m 1s |
|  | Kestrel | 8.93 | 1.28 | 2m 34s |
|  | Human | 10.15 | 3.18 | 6m |
|  | NCBI-v | 8.44 | 0.40 | 44s |
|  | SE | 11.15 | 1.25 | 1m 59s |
|  | HPRC | 11.50 | 5.35 | 9m 26s |
| 63 | Cod | 4.81 | 0.33 | 2m 22s |
|  | Kestrel | 4.28 | 0.62 | 4m 55s |
|  | Human | 5.30 | 1.83 | 12m 5s |
|  | NCBI-v | 4.37 | 0.23 | 1m 45s |
|  | SE | 7.59 | 1.45 | 6m 25s |
|  | HPRC | 7.76 | 5.75 | 25m 53s |

**Table 5.**
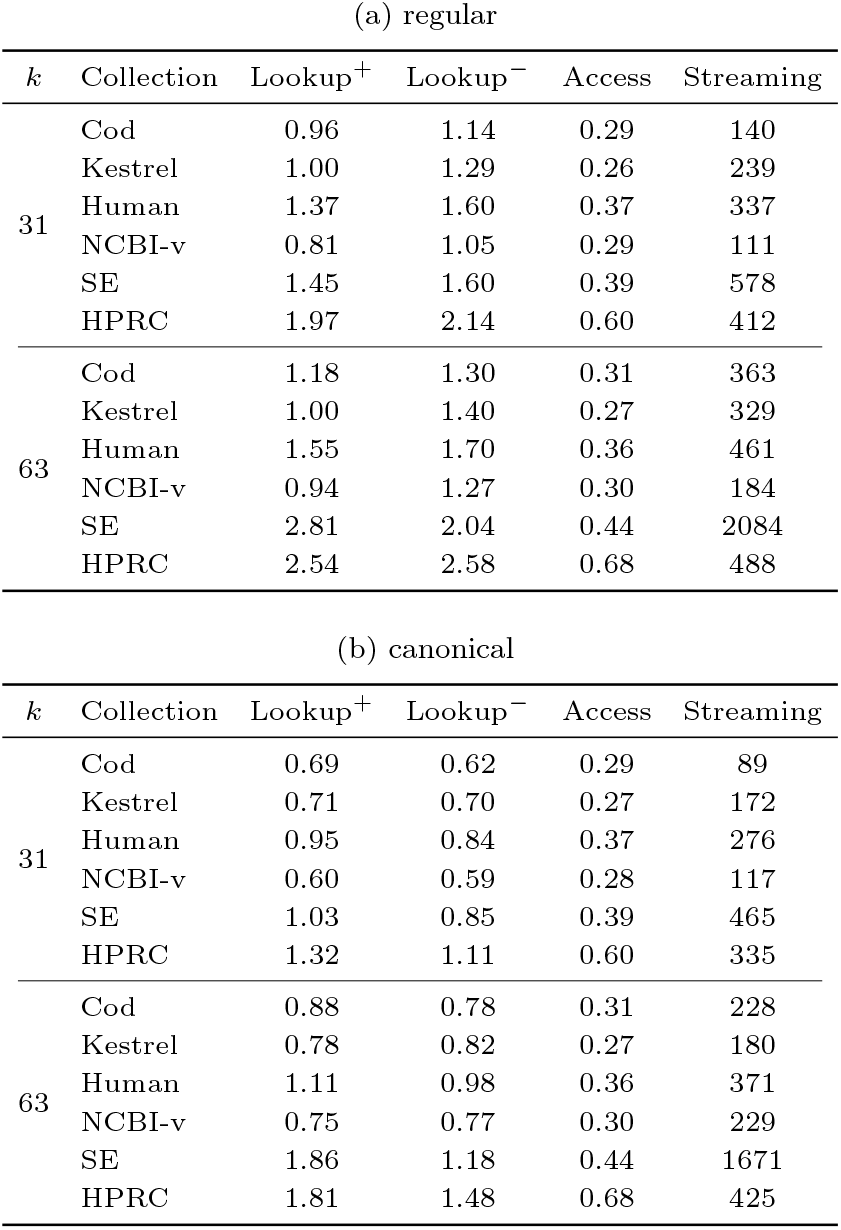
Query efficiency for previous SSHash. Timings for Lookup and Access are in (average) microseconds per k-mer. For Streaming, we report (average) nanoseconds per *k-mer*.

## Footnotes

1 An algorithm that computes 𝒮 such that both |𝒮| and N are minimum is known [Schmidt and Alanko, 2023], improving over previous heuristics [Rahman and Medvedev, 2020; Břinda et al., 2021]. In this case, the strings in 𝒮 are called *eulertigs*. This is the form of 𝒮 we are going to use in the experimental analysis in Section 8.

2 We omit ceiling operators when they are not essential.

3 Technically, there can be more than k *—* m +1 consecutive k-mers with the same minimizer. In this case, however, we can split the super-k-mer into chunks of at most k *—* m + 1 k-mers.

4 Technically speaking, we should choose l so that the number of minimizers with two occurrences is less than 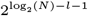. Due to the skew distribution of such occurrences for sufficiently long m, this is always the case for l = 6 across all of our experiments.

1 GitHub commit: `a2a2d26817fe3f476ceac44809c333ede6622ff3`.

